# Validation of known risk factors associated with carpal tunnel syndrome: A retrospective nationwide 11-year population-based cohort study in South Korea

**DOI:** 10.1101/253666

**Authors:** Hong-Jae Lee, Hyun Sun Lim, Hyoung Seop Kim

**Affiliations:** Department of Physical Medicine and Rehabilitation, Ilsanpaik Hospital, Inje University, Republic of Korea; Department of Policy Research Affairs, National Health Insurance Service, Ilsan Hospital; Department of Physical Medicine and Rehabilitation, National Health Insurance Service, Ilsan Hospital

**Author notes:** **Corresponding Author: Address:** National Health Insurance Service Ilsan Hospital, 100 Ilsan-ro Ilsan-donggu Gogyang Zip code: 10444 **Country:** Republic of Korea **E-mail:**.

**Keywords:** Carpal tunnel syndrome, Risk factors, Cohort Study, Korea

## Abstract

**Key Points:** *Question:* What is the relationship between the previously known risk factors and occurrence of carpal tunnel syndrome (CTS)?

**Findings:** In this retrospective population-based cohort study that included 512,942 participants sampled from the Korean National Health Insurance System database, we determined the following known risk factors were related to the occurrence of CTS: the age of 40s, female, being overweight, diabetes, rheumatoid arthritis, gout, and Raynaud’s syndrome. However, ESRD, hypothyroidism and smoking were not correlated with CTS occurrence.

**Implications:** We identified the age of 40s, female, overweight, diabetes, rheumatoid arthritis, gout, and Raynaud’s syndrome as risk factors for the occurrence of CTS.

**Abstract:** *Importance:* There have been few large-scale studies that have included a risk factor analysis for CTS. No prior study has investigated and validated the relationship between the occurrence of CTS and known risk factors using nationwide health care database.

*Objective:* To confirm the actual risk factors for CTS out of various known risk factors

*Design:* We conducted this study using a retrospective cohort model based on the combined two databases of the Korean National Health Insurance System; the national periodic health screening program database from 2002–2003 and health insurance database of reimbursement claims from 2003 through 2013.

*Setting:* A population-based retrospective cohort study.

*Participants:* First, we randomly sampled 514,795 patients who represented 10% of the 5,147,950 people who took part in periodic health screenings in 2002–2003. Existing CTS patients were excluded from this group. Therefore, this study finally included 512,942 participants and followed up their medical records from 2003–2013.

*Main Outcomes and Measures:* Desired outcomes were the incidence rate of CTS in patients with various risk factors and the hazard ratios of risk factors affecting the disease’s occurrence.

*Results:* The incidence of CTS was highest in patients in the age of 40s, in the moderate obesity group, in females, and in patients with diabetes mellitus (DM). The hazard ratio analysis revealed that the following risk factors were strongly related to the occurrence of CTS: age of 40s, female, obesity, DM, rheumatoid arthritis, gout, and Raynaud’s syndrome. However, ESRD, hypothyroidism and smoking were not correlated with CTS occurrence.

*Conclusions and Relevance:* In our large-scale cohort study, risk factors such as being in one’s 40s, obesity, being female, suffering from DM, and rheumatoid arthritis were reaffirmed as those of CTS occurrence.

## INTRODUCTION

Carpal tunnel syndrome (CTS) is the most common compressive neuropathy of the median nerve.^1-3^ This condition is thought to be caused by the entrapment of the median nerve within the tendons of the hand in the carpal tunnel. Clinical features include tingling sensations, numbness and neuropathic pain over the median nerve distribution, and thenar muscle weakness and atrophy. ^3^ The prevalence of CTS in the general population has been reported to be approximately 3.8–5.8%.^1,4,5^[1] The reason for this variation in the prevalence rate may be due to differences in diagnostic criteria, study designs and population.^2,3,6^ Although CTS has been studied extensively, its pathophysiology is still not fully understood.^7^ Several previous studies have revealed an association between various risk factors and CTS. For example, it is generally known to be common among middle-aged women.^2,4^ Other known risk factors are higher body mass index (BMI), diabetes mellitus (DM), rheumatoid arthritis (RA), gout, end-stage renal disease (ESRD), hypothyroidism, Raynaud’s syndrome (RS), occupation, trigger finger, computer use, acromegaly, and excessive alcohol abuse and smoking.^2,8-14^

The Korean health care system is based on the Korean National Health Insurance Service (KNHIS). Two independent institutions, the National Health Insurance Corporation (NHIC) and the Health Insurance Review and Assessment Service of Korea (HIRA), are the main offices that manage the medical insurance system. Both institutions compile data about health insurance operations and distribute them to researchers for study or to produce health policy. In South Korea, most people (97%) are required to enroll in the KNHIS.^3^ All medical institutions and facilities have mandatory contracts with the NHIC,^15^ which plays a main role in the qualification of insurance payers and beneficiaries, the imposition of premiums, and the prevention of diseases by operating programs such as national periodic health screening. The HIRA is in charge of reviewing reimbursement claims from clinics and hospitals.^15^ These claims are accompanied by data including diagnostic codes, personal information, and the direct medical costs of both inpatient and outpatient care and dental services.^15^ Thus, nearly all of the data in the health insurance system are centralized into one large database.

We obtained 11-year follow-up cohort data of the NHIC and HIRA and aimed to verify the risk factors that impact the condition’s occurrence.

## Methods

### Statement of Ethics

This study adhered to the tenets of the Declaration of Helsinki, and this research project was approved by the KNHIS (The research management number is NHIS-2017-2-536). This study design was reviewed and approved by the Institutional Review Board of the National Health Insurance Service, Ilsan Hospital, Gyeonggi-do, South Korea. Written informed consent was waived.

### Database

We used combined data from the national periodic health screening program database in 2002–2003 of the NHIC and the HIRA’s database of reimbursement claims from 2003 to 2013.^16,17^ The KNHIS and HIRA use the Korean Classification of Diseases (KCD) disease classification codes, which were modified from the International Classification of Diseases (ICD) codes.

### Study Sample

We compiled an 11-year follow-up cohort model by randomly sampling participants from the database. Ten percent of the total population who received periodic health screening in 2002 and 2003 were sampled, which produced 514,795 out of 5,147,950 total participants (Fig. 1). Among these, people who had already been diagnosed with CTS were excluded. Therefore, we began our study with 512,942 participants who did not have CTS at the beginning of the year.

**Fig 1.**
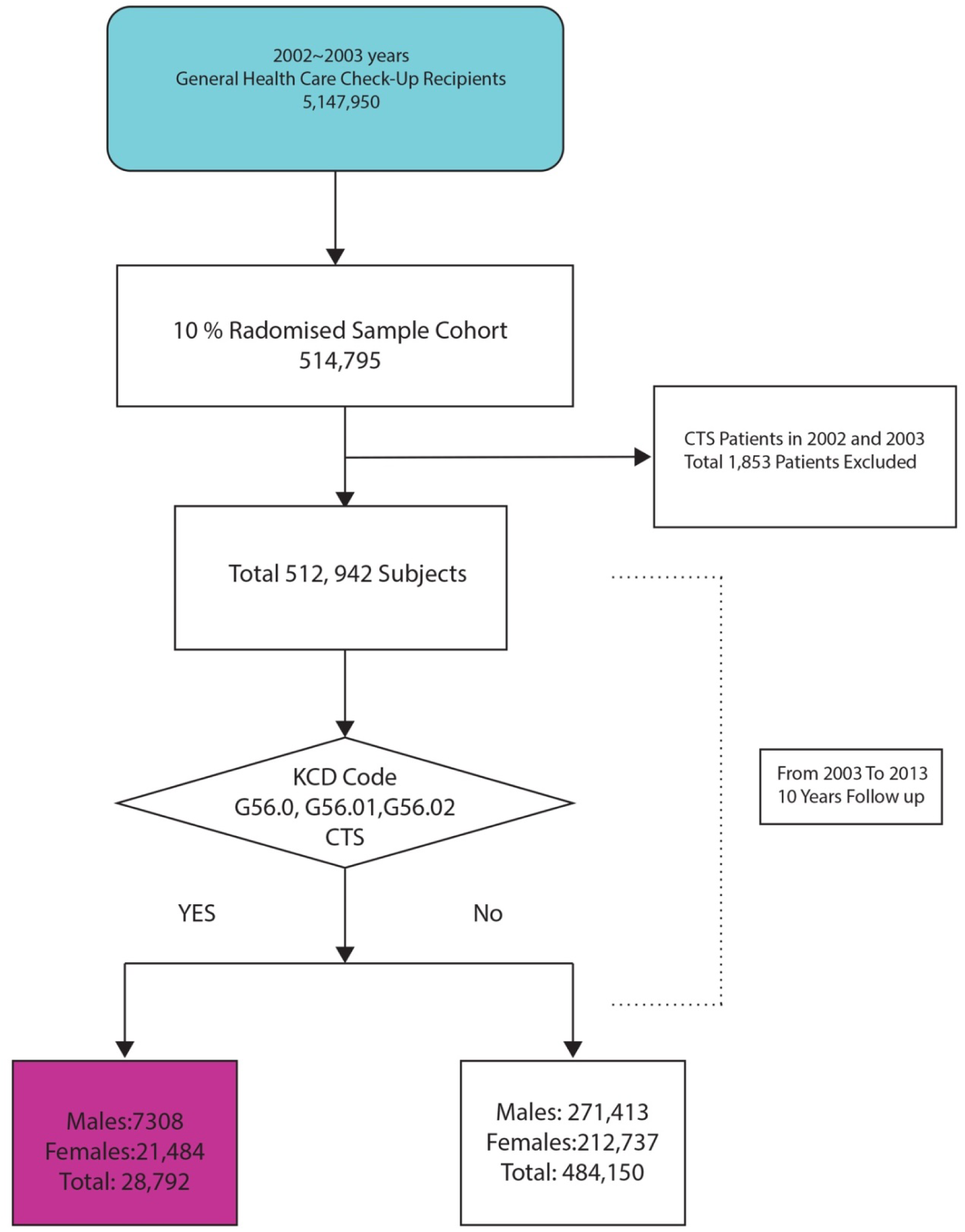
Flow chart for sampling and selecting the participants.

We defined CTS patients as those who had CTS codes as their main or secondary condition in the claims. We collected CTS patients who had the following diagnostic codes (KCD) based on data from the HIRA: carpal tunnel syndrome (G56.00), carpal tunnel syndrome in an unspecified upper limb (G56.01), carpal tunnel syndrome in the right upper limb (G56.02), or carpal tunnel syndrome in the left upper limb (G56.03).

### Validation of the known risk factors for carpal tunnel syndrome

The previously known risk factors were being female, being middle-aged, having a high BMI, smoking, and being diagnosed with hypertension (HTN), DM, RA, gout, ESRD, hypothyroidism, or RS. We validated the known risk factors for CTS by obtaining the incidence rate of CTS in patients who had each risk factor as well as the hazard ratio (HR) of each risk factor that leads the disease to occur.

BMI was classified into five grades based on the Asian standard as follows: below 18.5 (underweight), 18.5–22.9 (normal), 23–24.9 (overweight), 25–29.9 (moderate obesity), and 30–35 (severe obesity). Participant smoking history was classified as non-smoker, ex-smoker or current smoker. We defined the presence of comorbidities as the following diagnostic codes in claims for medical services: DM (E.10, E.11, E.12, E.13, E.14), RA (M.05, M.06), gout (M.10), ESRD (N.18), hypothyroidism (E.031, E.032, E.038, E.039), and Raynaud’s syndrome (I.730).

### Statistical analysis

We examined the occurrence rates of CTS according to various risk factors as well as hazard ratios that lead to CTS.

Descriptive statistics of the study populations were obtained, and Chi-square tests were performed to examine the association of risk factors with CTS. To identify any correlation between risk factors and CTS occurrence, adjusted hazard ratios (HRs) and 95% confidence intervals (CI) were determined using a multivariate Cox proportional hazard regression. A significance level of 0.05 was set. The statistical package SAS for Windows, version 9.2 (SAS Inc., Cary, NC, USA), was used to perform the analyses in this study.

## Results

### CTS and known risk factors

The incidence rate of CTS by risk factor is shown in Table 1. The overall incidence rate was 5.61% in adults over 40 years: 1.42% in males and 4.19% in females. The occurrence rates according to age group were 2.45% in their 40s, 1.86% in their 50s, 1.08% in their 60s, 0.22% in their 70s, and 0.11% in their 80s and older. The occurrence rates of CTS with diabetes mellitus and RA were 0.91% and 0.52%, respectively, which were higher than those for other comorbidities. Moderate obesity had the highest occurrence (2.10%) among the BMI groups. Interestingly, the non-smoker group had a much higher occurrence (4.66%) rather than the exsmoker or current smoker group.

**Table 1.** The incidence of carpal tunnel syndrome according to known risk factors.

|  | Carpal tunnel syndrome and its incidence |  |  |
| --- | --- | --- | --- |
|  | Yes | No | % |
| Overall | 28,792 | 484,150 | 5.61 |
| **Age** |  |  |  |
| 40s | 12,548 | 212,540 | 2.45 |
| 50s | 9,526 | 135,750 | 1.86 |
| 60s | 5,540 | 99,228 | 1.08 |
| 70s | 1,148 | 35,252 | 0.22 |
| 80s | 30 | 1,380 | 0.11 |
| **Gender** |  |  |  |
| Male | 7,308 | 27,1413 | 1.42 |
| Female | 21,484 | 212,737 | 4.19 |
| **Comorbidities** |  |  |  |
| Diabetes mellitus | 4,647 | 64,865 | 0.91 |
| Rheumatoid arthritis | 2,683 | 23,965 | 0.50 |
| Gout | 462 | 6,744 | 0.09 |
| End-stage renal disease | 91 | 1,371 | 0.02 |
| Hypothyroidism | 487 | 5,209 | 0.09 |
| Raynaud's syndrome | 56 | 305 | 0.01 |
| **BMI** |  |  |  |
| BMI ≤18.5 | 339 | 11,599 | 0.07 |
| 18.5< BMI ≤22.9 | 8,258 | 173,003 | 1.66 |
| 22.9< BMI ≤24.9 | 7,804 | 131,635 | 1.52 |
| 24.9< BMI ≤29.9 | 10,754 | 154,140 | 2.10 |
| 30< BMI | 1,339 | 13,322 | 0.26 |
| **Smoking History** |  |  |  |
| Non-smoker | 22,872 | 306,287 | 4.66 |
| Ex-smoker | 1,333 | 42,293 | 0.27 |
| Current smoker | 3,480 | 114,985 | 0.71 |

**Table 2.** Hazard ratios for known CTS risk factors.

| **Clinical Variables** | **HR** | **95% HR confidence limits** | **P-value** |
| --- | --- | --- | --- |
| **Age** | 0.977 | 0.971–0.974 | <.0001 |
| **Female** | 3.005 | 2.882–3.135 | <.0001 |
| **BMI £18.5** | 0.655 | 0.583–0.737 | <.0001 |
| **18.5< BMI £22.9** | 1 |  |  |
| **22.9< BMI £24.9** | 1.252 | 1.211–1.294 | <.0001 |
| **24.9< BMI £29.9** | 1.461 | 1.416–1.507 | <.0001 |
| **30< BMI** | 1.654 | 1.553–1.762 | <.0001 |
| **Non-smoker** | 1 |  |  |
| **Ex-smoker** | 1.044 | 0.980–1.113 | 0.1834 |
| **Current smoker** | 0.972 | 0.928–1.017 | 0.2170 |
| **DM** | 1.224 | 1.181–1.268 | <.0001 |
| **RA** | 1.448 | 1.385–1.515 | <.0001 |
| **Gout** | 1.267 | 1.145–1.402 | <.0001 |
| **ESRD** | 1.023 | 0.815–1.284 | 0.8428 |
| **Hypothyroidism** | 1.040 | 0.941–1.148 | 0.4440 |
| **RS** | 2.089 | 1.548–2.819 | <.0001 |

In our correlation analyses of known risk factors and the occurrence of CTS, the HR decreased as age increased (HR=0.977, p <0.0001); the HR of females was 3.005 (p <0.0001), which was almost three times higher than that of the male group. The HR was 1.252 (p <0.0001) in the overweight group, 1.461 (p <0.0001) in the moderate obesity group, and 1.654 (p <0.0001) in the severe obesity group. In addition, the HR was 1.224 for DM, 1.448 for RA, 1.267 for gout, and 2,089 for Raynaud’s syndrome with statistical significance (p <0.0001). However, ESRD, hypothyroidism and smoking were not correlated with CTS occurrence (Table 5).

## Discussion

In this study, we aimed to validate the known risk factors for CTS by obtaining the incidence rate of CTS in patients with each risk factor as well as the HR of each risk factor for the condition.

We used the NHIS database for the information needed to carry out this study.^17^ South Korea is a small country like Taiwan, and both nations have a National Health Insurance System. All health-related information has been collected into one central NHIS database, which makes it easy to set up a long-term cohort model. Researchers in Taiwan have already begun reporting the results of some studies that have used a nationwide cohort model based on this large database. This type of model is advantageous because it can enroll a large number of participants and allows researchers to carry out studies at low cost and with little manpower, which may lead to influential new results or the verification of existing research results that were previously obtained from small populations.

### A. Validation of known risk factors related to CTS occurrence

The basic mechanism of CTS’s occurrence is ischemia of the median nerve caused by increased pressure within the carpal tunnel due to a disorder of the intraneural microcirculation of the nerve or by the alteration of the surrounding connective tissue, such as subsynovial connective tissue hypertrophy and synovial tissue hypertrophy.^18-20^

This study showed that risk factors including female, age of 40s, a higher BMI, and a diagnosis of DM, RA, gout, or RS were related to the onset of CTS. It is well known that CTS is more common in women, especially in middle age. The occurrence of CTS in women is two to four times higher than in men.^6,21^. Our study also showed that the occurrence ratio of females to males was 2.95.

Many studies have reported that increasing age is associated with the onset of CTS.^21-23^ However, in our study, the HR for age was 0.997, which indicated that the occurrence of CTS decreased as age increased. One possible explanation for this is that the population in the current study was limited to participants who received periodic health screenings and were in their 40s or older. Considering that the highest occurrence was in their 40s (2.45%), we can also infer that overuse of the hands and wrists during this period may be one of the causative factors of CTS. During their 40s, many women are still working or are busy taking care of babies and doing household chores, which may increase their risk.

Our study showed that a higher BMI was associated with a higher HR. These findings coincided with those of previous studies that reported higher BMI as one of the major risk factors for CTS.^21,23-26^ The suggested hypothesis is that increased fat tissue inside the carpal tunnel increases hydrostatic pressure or that water accumulation is accelerated in connective tissues and therefore causes median nerve compression.^7,23,24,26^

DM is also a well-known risk factor for CTS.^23,27,28^ Our results supported this notion because the HR of DM was >1. Previous studies have reported that CTS is involved in up to one-third of patients with DM and is three times as prevalent in a diabetic population compared to a healthy population.^27,29^ The reasons for this difference are that the median nerve undergoes repeated yet undetected microtrauma and fluid accumulates within the confined space of the carpal tunnel due to metabolic changes.^29,30^ A nationwide population-based cohort study conducted in Taiwan on patients with DM revealed that women and younger patients with DM had the highest risk for diabetic hand syndromes (DHS), including CTS. ^31^ However, a different study reported that type II DM was not a risk factor for CTS.^32^ Although it can be argued that it is difficult to clarify the pathogenesis of CTS in diabetic patients,^28^ our results support the claim that DM is a major risk factor for CTS.

RA is a chronic inflammatory disease accompanied by various extra-articular manifestations and progressive articular damage.^33^ It frequently causes tenosynovitis and can anatomically alter the carpal tunnel.^34^ RA is a known risk factor for CTS because the tenosynovitis caused by RA increases intracarpal pressure and injures the median nerve.^29,34,35^ The result of our study supported that RA is one risk factor for CTS.

Similarly, gout is also known to cause inflammation in soft tissues and to produce gouty tophi, which can cause CTS.^36,37^ In previous studies, gout was reported to be a comorbidity associated with CTS,^36,38^ and this claim is consistent with our findings.

RS is caused by vasculitis associated with systemic inflammatory disorders and results in impaired microcirculation.^39,40^ Autonomic dysfunction may occur in RS or CTS, which can produce Raynaud’s phenomenon symptoms^41^. Alternatively, both conditions may be present concurrently.^39,40^ A meta-analysis found that CTS and RS were statistically related. In our study, the HR of RS (2.089) was the second highest among all CTS risk factors.

According to our results, ESRD, hypothyroidism and smoking were not associated with CTS. Previous studies have identified a controversy between smoking and CTS. One meta-analysis reported that current smoking and CTS were associated in cross-sectional studies but not in cohort studies,^42^ although one cross-sectional study with a population of 514 found that smoking decreased the incidence of CTS.^27^ However, men are more likely to be smokers than women, so smoking may be a compounding factor; therefore, cross-sectional studies are insufficient to conclude that there is a correlation between smoking and CTS.^42^

ESRD patients currently receiving renal dialysis are likely to develop carpal tunnel syndrome due to amyloid deposits in the soft tissue that are similar to gout tophi. In fact, many previous studies have reported a correlation between ESRD and the occurrence of CTS.^43-45^ However, most of these studies were conducted using small populations;^44,45^ large-scale studies have not yet been conducted. Our investigation of a large population revealed that ESRD was not associated with CTS. Thus, we think the results of our study are more objective.

In one cross-sectional study, a correlation was identified between hypothyroidism and CTS^27^. In a different meta-analysis, there was a modest association between hypothyroidism and CTS, but evidence of a publication bias may account for this correlation.^46^ An investigation of 1 million people in Taiwan reported that CTS that occurred in patients under the age of 39 was related to hypothyroidism, but CTS in those over 40 was not.^38^ The results of our study conducted using large population were in agreement.

A strong point of our study is that it was a nationwide, 11-year follow-up cohort model with sample size over 500,000; the quality of the data is also objective and reliable because we used the NHIS database. A second merit of our study is that we carried out several research tasks at once: validation of the known risk factors for CTS.

Our study also had a few limitations. First, we did not include any participants under the age of 40 years because the national periodic health screening is only offered to people in their 40s or older. Second, the diagnosis of CTS was made using the results of electrodiagnostics (electromyelography and nerve conduction velocity) along with the clinical impression of doctors who examined the patients. In addition, this study assumed that the risk factors that existed during the initial year from 2002–2003 would persist for the next 10 years unchanged or with a little change. However, smoking and BMI are subject to change.

## Conclusion

In our study, we validated the following risk factors for CTS: female, an age from 40–49 years, high BMI, DM, gout, and RS. However, smoking, hypothyroidism and ESRD were not associated with CTS. We believe that the results of our study will be helpful in determining the pathophysiology of CTS.

